# Serotonergic control of spine morphology during growth in zebrafish

**DOI:** 10.64898/2026.09.20.753056

**Authors:** Stevana R. Schauer, Bryson Tyler B. Ricamona, Maximilian R. Segeberg, Laura Desban, Zane L. Walsh, Judith S. Eisen, Daniel T. Grimes

**Affiliations:** Institute of Molecular Biology, Department of Biology, University of Oregon Eugene, OR United States; Institute of Neuroscience, Department of Biology, University of Oregon Eugene, OR United States

**Keywords:** serotonin, scoliosis, zebrafish, vertebral morphology, pineal gland, neuromuscular

## Abstract

Scoliosis is a three-dimensional deformity of the vertebral column that commonly emerges during growth, yet the biological mechanisms that initiate and drive curve progression remain poorly understood. Serotonin (5-hydroxytryptamine [5-HT]) has long been linked to scoliosis through its relationship with pineal melatonin signaling and through genetic associations with human disease, but its role in spinal morphogenesis has remained unclear. Here, using zebrafish, we show that serotonergic signaling is required for normal spine morphology. Transient exposure to 5-HT during development induces later spinal abnormalities, while loss of Tph2, an enzyme required for neuronal 5-HT synthesis, causes progressive three-dimensional spinal curvature during juvenile growth. *tph2* mutants nevertheless exhibit initially normal vertebral patterning, arguing against an early skeletal defect as the initiating cause of curvature. Additionally, sustained ablation of *exorh*-positive pineal photoreceptors does not induce spinal curvature, arguing against a pineal origin for the phenotype. The Reissner fiber, which has also been linked to spinal curvature in zebrafish, formed normally in *tph2* mutants. Instead, mutants showed a marked reduction in Fev-expressing intraspinal serotonergic neurons together with increased locomotor activity, characterized by longer swim bouts and shorter periods of inactivity between bouts. Importantly, these neuronal and motor defects were evident before the onset of overt spinal curvature. Together, these findings identify neuronal serotonin as a critical regulator of spinal stability and support a model in which altered serotonergic control of motor function changes the mechanical environment experienced by the growing spine, increasing its susceptibility to progressive curvature.

## Introduction

During growth, the vertebral column must maintain its overall shape while both elongating and experiencing changing mechanical forces from surrounding tissues and the wider environment, including gravity, muscle contraction, and movement. A breakdown of spinal morphology occurs in scoliosis, when aberrant three-dimensional curvature of the vertebral column develops. In the most common form of scoliosis, adolescent idiopathic scoliosis (AIS), otherwise healthy children develop >10° lateral spinal curves during periods of adolescent growth. Though a wide range of biological and physiological differences have been associated with AIS, spanning skeletal, muscular, neurological, and sensorimotor systems, it has been challenging to understand the underlying causes of spinal curvature (Cheng et al., 2015; Wajchenberg et al., 2016; Peng et al., 2020). Notably, a major difficulty is separating changes that drive the initiation of scoliosis from those that develop secondarily in response to an already curved spine.

Vertebrate animal models therefore provide powerful tools for testing how spinal morphology is generated and maintained, and for distinguishing causes of curvature from its consequences. In particular, zebrafish (*Danio rerio*) are especially tractable models because spinal morphology can be followed throughout growth while genetic perturbations can produce progressive spinal curve phenotypes that resemble key features of AIS (Buchan et al., 2014; Hayes et al., 2014; Grimes et al., 2016; Boswell and Ciruna, 2017; Bearce and Grimes, 2021; Gray et al., 2021). Studies in zebrafish have revealed that normal axial morphology depends on processes that act well before overt spinal curvature manifests. For instance, motile cilia-generated cerebrospinal fluid (CSF) flow in the central nervous system (CNS) is essential for maintaining a well-aligned spine (Grimes et al., 2016; Konjikusic et al., 2018; Wang et al., 2020; Wang et al., 2022; Xie et al., 2023). A prominent role of CSF flow is the formation of the Reissner fiber (RF), a long extracellular proteinaceous thread that forms in the brain ventricles and along the spinal canal (Reissner, 1860; Cantaut-Belarif et al., 2018; Grimes, 2019; Sepúlveda et al., 2021). Disruption of RF formation or maintenance causes axial curvature and scoliosis-like phenotypes in zebrafish (Cantaut-Belarif et al., 2018; Lu et al., 2020; Rose et al., 2020; Troutwine et al., 2020). When the cilia-CSF-RF axis is disrupted, spinal curves develop during growth without major vertebral structural malformations, similar to the situation in AIS patients.

Animal models have also suggested that the pineal gland and its secreted product melatonin may play a role in spinal alignment. Beginning with classical experiments in the 1950s, surgical removal of the pineal gland has been reported to induce scoliosis-like curves in several vertebrate species (Pflugfelder, 1953; Thillard, 1959; Machida et al., 1993; Kanemura et al., 1997; O’Kelly et al., 1999; Cheung et al., 2005; Man et al., 2014). Human genetic studies have also reported associations between components of melatonin signaling and AIS (Qiu et al., 2006). These findings led to the long-standing hypothesis that pineal and/or melatonin dysfunction contributes to scoliosis (Machida et al., 1995, 1996; Bagnall et al., 1996; Wang et al., 1998; Cheung et al., 2003; Turgut et al., 2003; Moreau et al., 2004; Girardo et al., 2011). However, this idea remains controversial primarily because pinealectomy and melatonin reduction do not consistently produce curvature across experimental systems and because the curves that are induced are dissimilar in some respects to those exhibited by AIS patients (Hilibrand et al., 1996; O’Kelly et al., 1999; Cheung et al., 2003; Cheung et al., 2005; Day et al., 2007; Fagan et al., 2009). Interpretation of these experiments is additionally complicated by the invasiveness of surgical pineal removal and the possibility of damage to surrounding tissues, including the subcommissural organ (SCO), which secretes the major protein component of the RF, SCO-spondin (Sspo; Grondona et al., 2012), and other potentially important neuronal populations.

A growing body of evidence indeed points to neural control of posture as an important contributor to spinal stability. It has been suggested that maintaining an upright vertebral column requires continuous integration of vestibular and proprioceptive information with motor outputs that regulate axial and paraspinal muscle activity. Abnormal sensory integration could therefore produce an altered internal representation of verticality, leading to chronically abnormal postural motor output (Hawasli et al., 2015). In agreement, differences in postural control, balance, vestibular processing, and paraspinal muscle structure have all been described in individuals with AIS (Herman et al., 1985; Catanzariti et al., 2014; Dufvenberg et al., 2018; Kim et al., 2020; Lau et al., 2022; Chan et al., 2024; Paramento et al., 2024). Locomotion provides a related neuromuscular route through which the nervous system might influence spinal morphology, because each bout of movement recruits axial musculature and imposes mechanical forces on the vertebral column. The frequency, duration, symmetry, and magnitude of muscle activity therefore determine, in part, the mechanical environment in which the spine grows. Perturbation of this environment may alter spinal alignment (Stokes et al., 1996; Stokes, 2007). Animal studies further support the idea that neural and neuromuscular perturbations can influence spinal morphology (Zhang et al., 2018; Vesque et al., 2019; Lu et al., 2020; Bearce et al., 2022; Gaillard et al., 2023; Endo et al., 2024; Wang et al., 2024, 2025; O’Hara-Smith et al., 2026), although the mechanisms linking altered neural activity to curve initiation remain poorly understood.

The neurotransmitter serotonin (5-hydroxytryptamine; 5-HT) has emerged as a candidate regulator of spinal morphology. Genetic studies demonstrate associations between variants in *TPH1*, which encodes a 5-HT synthesis enzyme, and susceptibility to AIS, as well as continued curve progression during brace treatment (Wang et al., 2008; Xu et al., 2011). These associations have not been consistently replicated across populations, however, making their biological significance uncertain (Takahashi et al., 2011; Li et al., 2021). Nevertheless, 5-HT is a particularly intriguing candidate because it plays roles in several biological systems that have independently been implicated in scoliosis.

First, 5-HT is the immediate biosynthetic precursor of melatonin and also acts as an autocrine signal that potentiates melatonin synthesis (Lee et al., 2021). Second, 5-HT signaling could interact with the cilia-CSF-RF pathway since the SCO receives serotonergic innervation while experimental manipulation of 5-HT signaling alters SCO secretory activity and Sspo expression (Bouchaud, 1979; Léger et al., 1983; Sakumoto et al., 1984; Richter et al., 2004). Third, 5-HT is a major modulator of spinal motor circuits controlling locomotion and axial muscle activity (Brustein et al., 2003; Gabriel et al., 2009; Montgomery et al., 2018). Thus, altered serotonergic signaling could conceivably influence spinal morphology through one or more of these mechanisms. However, direct evidence that serotonergic signaling itself plays a causal role in maintaining spinal morphology is lacking.

Here, we used zebrafish to address this question by determining whether serotonergic signaling is required to maintain spine morphology, and to investigate mechanisms through which it might play such a role. We show that loss of Tph2, an enzyme responsible for neuronal 5-HT synthesis, leads to the emergence of three-dimensional spinal curvature during growth. Curves appear despite initially normal vertebral patterning and formation, similar to AIS. To test whether pineal function may be involved in Tph2-dependent phenotypes, and avoid the potentially confounding effects of surgical pinealectomy, we used targeted, genetically encoded chemical ablation of the pineal gland. However, this had no effect on spine morphology. Similarly, the RF remained intact in *tph2* mutants across multiple stages. Instead, mutants exhibited a reduction in serotonergic intraspinal neurons (ISNs) and increased locomotor activity. Importantly, these phenotypes were present before the onset of overt spinal curvature. These findings identify serotonergic signaling as a regulator of spinal morphology during growth and suggest that altered neural and neuromuscular function may change the mechanical environment experienced by the developing vertebral column, making it susceptible to curvature.

## Results

### The 5-HT synthesis enzyme, Tph2, is required for normal zebrafish spine morphology

To determine whether serotonergic signaling can influence spine morphology, we first exposed wild-type (WT) zebrafish larvae to exogenous 5-HT by adding 1 or 2 mM 5-HT to the rearing water from the 1-cell stage until 6 days post fertilization (dpf). Animals were then raised to 2.5 months post fertilization (mpf) for skeletal analysis by X-ray imaging. Mock-treated zebrafish exhibited normal spine morphology and well-formed vertebrae. In contrast, fish treated with 5-HT showed spinal curves and apparent vertebral defects at both concentrations tested (Fig 1A; 5/16 for 1 mM 5-HT and 4/15 for 2 mM 5-HT), in addition to craniofacial abnormalities (S1A Fig), consistent with the established role of serotonergic signaling in craniofacial development (Reisoli et al., 2010). Thus, transient exposure to 5-HT during embryonic and larval stages can disrupt later vertebral and spinal morphology.

**Fig 1.**
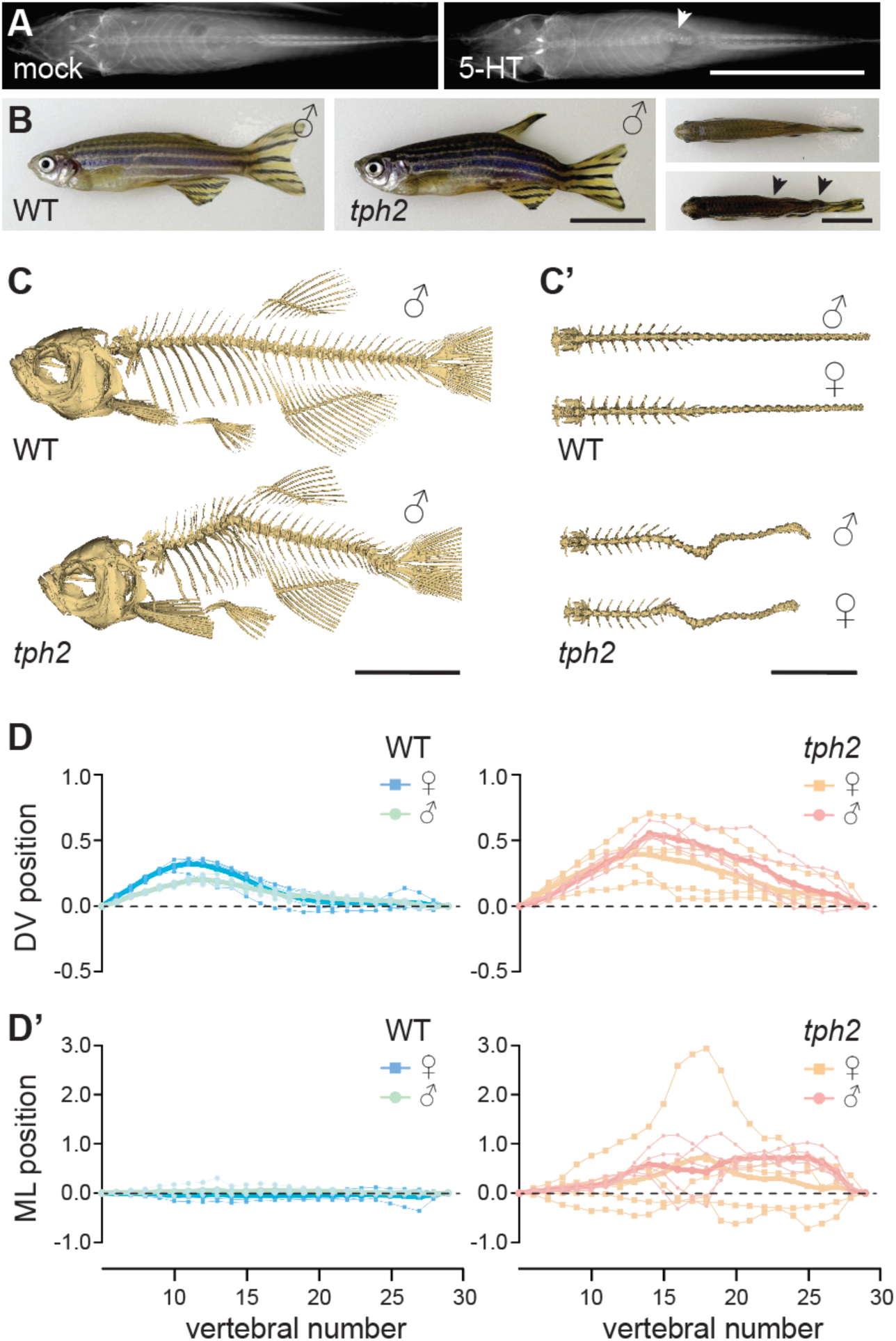
*tph2* mutants exhibit abnormal spine morphology. **(A)** 2.5 mpf WT fish either mock-treated or treated with 2 mM 5-HT from 0–6 dpf. Treated fish developed subtle spinal curvature (arrowhead). Scale bar = 10 mm. **(B)** Adult WT and *tph2* mutant males at 6 mpf shown in lateral and dorsal views. Arrows mark the positions of lateral spinal curvatures in *tph2* mutants. Scale bar = 10 mm. **(C–C’**) Lateral (C) and dorsal (Cʹ) views of three-dimensional reconstructions from *µ*CT scans of WT and *tph2* mutants at 3 mpf. Scale bar = 10 mm. **(D–D’)** Quantification of spinal shape from aligned vertebral coordinates in WT and *tph2* mutants at 3 mpf. Dorsoventral (DV) and mediolateral (ML) vertebral positions are plotted along the length of the spine. Positions are shown in arbitrary units and represent relative displacement of each vertebra from the reference spinal axis. Each line represents an individual fish; females and males are indicated separately. Thicker lines show means for each sex. *n* = 10 fish per genotype.

Next, we tested the consequences of reduced serotonergic signaling. Tryptophan hydroxylase 2 (Tph2) is expressed throughout the zebrafish CNS including the spinal cord, and is the principal enzyme required for neuronal 5-HT synthesis (Bellipanni et al., 2002; Walther et al., 2003; Montgomery et al., 2016). We therefore asked whether reducing neuronal 5-HT synthesis through genetic loss of Tph2 is sufficient to disrupt spinal morphology. We raised homozygous *tph2* mutants to adulthood and found they exhibited conspicuous axial deformities, including prominent kinks near the dorsal and caudal fins (Fig 1B; Oikonomou et al., 2019). Three-dimensional reconstructions of micro-computed tomography (*µ*CT; Bearce et al., 2023) scans revealed substantial spinal curves in *tph2* mutants (Fig 1C–1Cʹ; S1B and S1C Fig; S1 and S2 Videos). To quantify spinal shape, we compared the dorsoventral (DV) and mediolateral (ML) positions of successive vertebrae along the spine (Voigt et al., 2026). In the DV plane, WT fish showed the expected smooth and shallow dorsal curve (Fig 1C). By contrast, *tph2* mutants showed markedly greater DV displacement across the vertebral column (Fig 1D; S1B and S1C Fig; *p* = 4.72 × 10^-4^, permutation test), especially in vertebrae 12–22, encompassing the posterior trunk and extending into the caudal vertebral column. In the ML direction, curves occurred in either direction, with no left or right bias. As such, we compared absolute ML displacement and found substantially greater lateral displacement in *tph2* mutants compared with WT fish (Fig 1Cʹ and 1Dʹ; S1B and S1C Fig; *p* = 3.15 × 10^-5^, permutation test). Compared with the DV phenotype, the location and direction of ML displacement were more variable among individual mutants, although the largest deviations from WT occurred predominantly in the mid-to-posterior vertebral column (Fig 1Cʹ and 1Dʹ; S1B and S1C Fig).

Because AIS shows a pronounced female bias in curve prevalence and progression (Konieczny et al., 2013), and sex-dependent effects have also been observed in zebrafish scoliosis models (Buchan et al., 2014; Marie-Hardy et al., 2021), we next compared spinal curvature between male and female *tph2* mutants. However, we detected no evidence of sex-dependent differences in either DV (*p* = 0.254) or ML curvature (*p* = 0.889, permutation tests), indicating that the phenotype affects both sexes similarly. Overall, these findings demonstrate that Tph2 is required for normal spinal alignment in zebrafish, supporting a causal role for reduced neuronal 5-HT synthesis in the development of spinal curvature.

### Spinal curvature appears during juvenile growth and precedes vertebral malformations in *tph2* mutants

Next, we monitored *tph2* mutants and WT siblings through growth to determine when spinal curvature first emerges. Mutants grew at a similar rate to WT animals (S2A Fig; *p* = 0.936, two-way ANOVA). Spinal curves first became apparent at 21 dpf in a small proportion of mutants and subsequently increased in penetrance during juvenile growth (Fig 2A–2B). By 91 dpf, 71% of *tph2* mutants exhibited conspicuous spinal curves (Fig 2B), with penetrance being 100% when assessed at 1 year of age (*n* > 100). Notably, ML curvature generally preceded the appearance of DV curvature. These data indicate that spinal curvature arises at variable times during juvenile growth in *tph2* mutants and that the three-dimensional phenotype develops progressively, rather than appearing simultaneously in the DV and ML planes.

**Fig 2.**
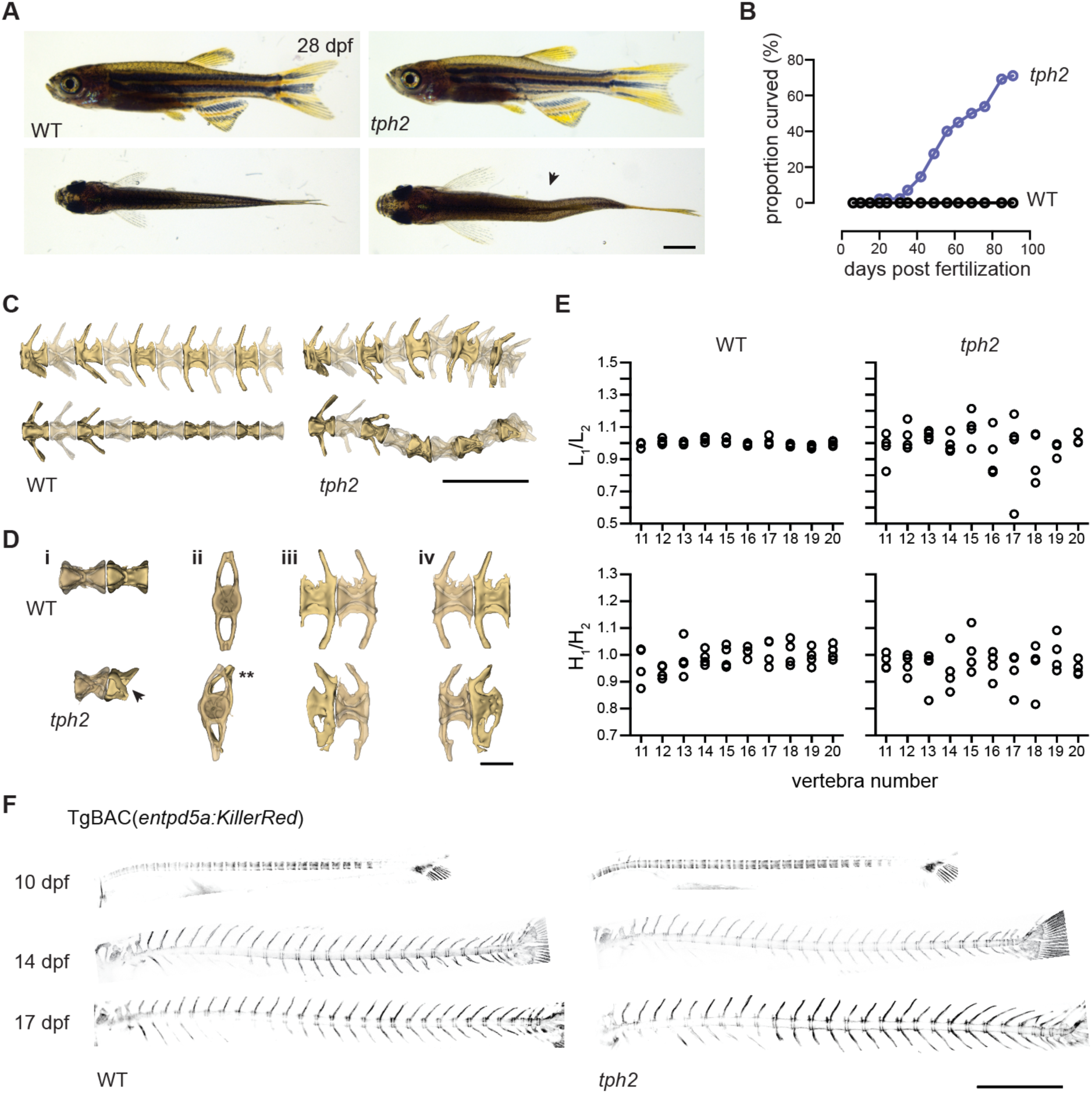
Spinal curvature in *tph2* mutants arises during juvenile growth after normal vertebral patterning. **(A)** Representative lateral and dorsal bright-field images of WT and *tph2* mutant zebrafish at 28 dpf. The arrowhead marks a spinal curve in the *tph2* mutant. Scale bar = 1 mm. **(B)** Developmental onset of spinal curvature. Fish were examined during juvenile growth and scored for the presence of overt spinal curvature. Curves first became apparent at 21 dpf in some fish then increased in penetrance during subsequent growth. **(C)** Representative three-dimensional reconstructions of isolated 6 mpf WT and *tph2* mutant spines generated from *µ*CT datasets, shown in lateral (top) and dorsal (bottom) views. *tph2* mutant spines show three-dimensional curvature and vertebral distortion. Scale bar = 1 mm. **(D)** Representative vertebrae from 6 mpf WT and *tph2* mutant animals illustrating abnormalities associated with curved regions. Compared with WT vertebrae, mutant vertebrae show wedge shapes (i), twisting (ii), compression (iii), and occasional fusion between adjacent vertebrae (iv). Scale bar = 250 *µ*m. **(E)** Quantification of vertebral morphology across vertebrae 11–20 in WT and *tph2* mutants. Ratios of longitudinal dimensions (L_1_/L_2_; top) and DV dimensions (H_1_/H_2_; bottom) are plotted for individual vertebrae. Vertebrae from *tph2* mutants show increased morphological variability, consistent with local vertebral distortion. **(F)** Representative images of TgBAC(*entpd5a:KillerRed*) in WT and *tph2* mutants at 10, 14 and 17 dpf. Mineralizing domains of the notochord sheath show regular segmentation in both *tph2* mutants and controls before overt spinal curvature. Scale bar = 1 mm.

In congenital scoliosis, curvature results from primary defects in vertebral formation or segmentation (Giampietro et al., 2009). By contrast, AIS develops in an initially normally patterned spine, although vertebral deformation can emerge secondarily as curves progress (Cheng et al., 2015). We therefore asked whether spinal curvature in *tph2* mutants originates from abnormal vertebral development or instead emerges after normal vertebral formation. In adult *tph2* mutants, three-dimensional *µ*CT reconstructions revealed localized vertebral abnormalities within regions of curvature (Fig 2C and 2D; S3 and S4 Videos). Individual vertebrae were variably compressed, warped, rotated, and, rarely, appeared fused in some severely affected adults (Fig 2C and 2D; S3 and S4 Videos). Indeed, quantitative analysis of isolated vertebrae from positions 11–20 along the spine revealed alterations in vertebral morphology within curved regions of *tph2* mutants (Fig 2E). Vertebral centrum shape was assessed by measuring rostrocaudal length along the dorsal and ventral surfaces and DV height at the anterior and posterior ends of each vertebral body then calculating length (L_1_/L_2_) and height (H_1_/H_2_) aspect ratios (Bearce et al., 2022). Values close to 1 therefore indicate a symmetric vertebra, whereas increasing deviation from 1 reflects greater vertebral distortion. Compared with WT vertebrae, *tph2* mutants showed a significantly greater mean displacement of the length aspect ratio from 1 (*p* = 0.014), whereas displacement of the height aspect ratio was not significantly different (*p* = 0.292, Welch’s *t* test), possibly due to increased variability in this metric in controls. Thus, adult *tph2* mutants exhibit modest vertebral abnormalities localized to curved regions of the spine.

These abnormalities could arise early during vertebral development and subsequently contribute to spinal curvature, or instead develop secondarily as the spine deforms. To distinguish these possibilities, we examined vertebral development before and during curve onset. We crossed *tph2* mutants to TgBAC(*entpd5a:KillerRed*), which labels mineralizing domains of the notochord sheath that subsequently give rise to vertebral centra (Wopat et al., 2018). This transgene therefore provides a marker of early vertebral patterning and centrum formation before vertebrae mature. In zebrafish, segmented *entpd5a*+ domains appear from around 4 dpf, showing the position of future chordacentra. Centrum mineralization then begins around 6 dpf and progresses over the following days and weeks (Xi et al., 2013; Wopat et al., 2018; Bagnat and Gray, 2020). We counted the number of *entpd5a+* centra and found the number in *tph2* mutants to be not different to WT controls at 10 dpf (*p* = 1.00), 14 dpf (*p =* 0.73), and 17 dpf (*p* = 0.46, Mann-Whitney tests). Moreover, centrum and vertebral shape were unchanged (Fig 2F). Together, these data indicate that early vertebral patterning and centrum formation proceed normally in *tph2* mutants through the period preceding curve onset, suggesting that the vertebral abnormalities observed in adults arise later.

We next examined vertebral morphology immediately after curves first became apparent using high-resolution *µ*CT. Vertebrae from heterozygous siblings were morphologically normal, whereas *tph2* mutants exhibited localized vertebral warping at the curve apex (S5 Video). Importantly, these animals did not yet show the severe defects, such as fusions, that were present in some adults. Small mineralized fragments were occasionally visible between distorted vertebrae (S2B Fig), consistent with early structural remodeling or damage accompanying the developing curve.

Overall, these findings indicate that *tph2* mutants undergo initially normal vertebral patterning before developing spinal curvature during juvenile growth. Local vertebral distortion appears as the curves emerge, while more severe abnormalities, including occasional vertebral fusions, develop later within chronically curved regions. The timing of curve onset and subsequent vertebral remodeling parallel AIS, in which spinal curvature emerges during growth and vertebral wedging and rotation can develop progressively with the curve (Cheng et al., 2015).

### Pineal photoreceptor ablation does not induce spinal curvature, and the RF remains intact in *tph2* mutants

Next, we asked which serotonergic populations are responsible for maintaining spine morphology. Given the longstanding hypothesis that the pineal gland and melatonin are needed for spinal alignment, and since *tph2* is expressed in the zebrafish pineal (Teraoka et al., 2004), we reasoned that Tph2 may play a pineal-dependent role in spine morphology. To test this, we ablated the pineal photoreceptor cells for the first 3.5 months of life using targeted and precise chemical ablation. We generated Tg(*exorh:mCherry-P2A-NTR2.0*) transgenic zebrafish in which the *exorh* promoter drives both mCherry and the enhanced nitroreductase (NTR) 2.0 (Asaoka et al., 2002; Sharrock et al., 2022). *exorh* is expressed in rod-like pineal photoreceptors, the predominant photoreceptor population in the pineal that is required for melatonin production (Asaoka et al., 2002; Anasarias-Dumo et al., 2026; Wexler et al., 2026). NTR 2.0 converts the prodrug metronidazole (MTZ) into a cytotoxic metabolite, allowing selective ablation of NTR-expressing cells. Transgene expression was restricted to the pineal gland (Fig 3A–3B; S3 Fig). Transgenic animals were exposed to MTZ continuously from the 1-cell stage to 4 dpf and subsequently retreated for 24 h every 3 days to prevent pinealocyte regeneration. This treatment regimen robustly ablated the pineal photoreceptors. Loss of pineal fluorescence and 5-HT immunoreactivity was confirmed throughout the experiment by immunostaining at early, intermediate, and final time points (Fig 3A–3B; S3 Fig).

**Fig 3.**
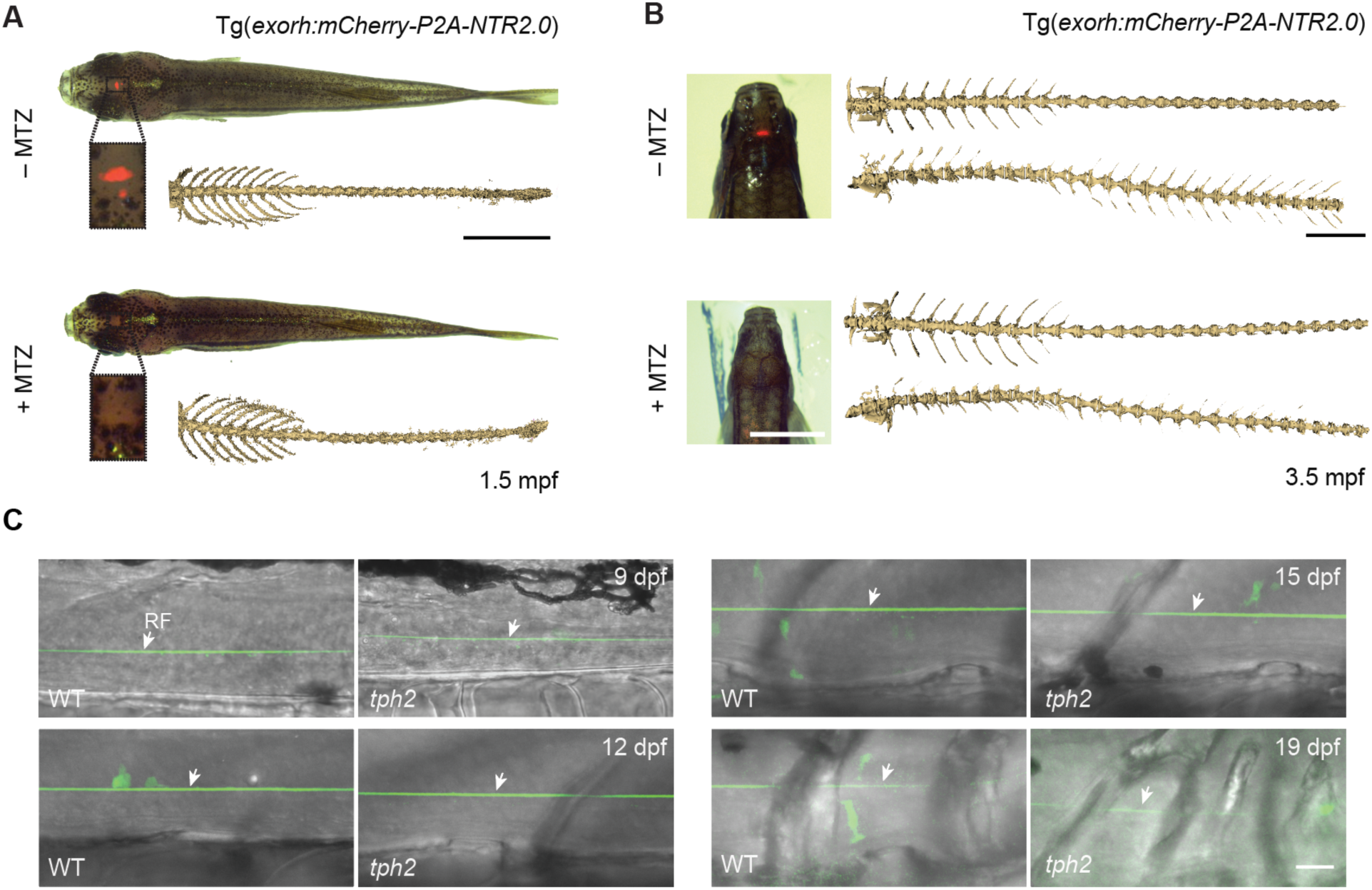
Pineal ablation does not induce spinal curvature and the Reissner fiber remains intact in *tph2* mutants. **(A)** Representative dorsal views of Tg(*exorh:mCherry-P2A-NTR2.0*) animals at 1.5 mpf with or without MTZ treatment. Insets show the pineal region. *µ*CT reconstructions are shown alongside each fish. Scale bar = 0.5 mm. **(B)** Representative dorsal views of Tg(*exorh:mCherry-P2A-NTR2.0*) animals at 3.5 mpf with or without MTZ treatment. *µ*CT reconstructions show dorsal (top) and lateral (bottom) views of the spine. The ribs have been removed. Scale bar = 0.5 mm. **(C)** Representative images of the RF in 9 dpf (WT: *n* = 2; *tph2: n* = 2), 12 dpf (WT: *n* = 4; *tph2: n* = 3), 15 dpf (WT: *n* = 9; *tph2: n* = 2), and 19 dpf (WT: *n* = 6; *tph2: n* = 4) fish. Arrows point to intact fibers. Scale bar = 20 *µ*m.

We then followed pineal photoreceptor-ablated animals through the period during which spinal curvature emerges in *tph2* mutants. At 1.5 mpf, none of the MTZ-treated NTR-positive animals exhibited spinal curvature (0 of 47; Fig 3A). In an independent cohort followed to 3.5 mpf, spinal curvature was again absent from all pineal-ablated animals (0 of 23; Fig 3B). Together, these results show that sustained loss of the pineal gland is not sufficient to induce scoliosis in zebrafish.

It is possible that the induction of spinal curves in some pinealectomized animals was caused by damage to the SCO and thereby disruption of the RF, a structure which is required for normal spinal morphology in zebrafish (Lu et al., 2020; Rose et al., 2020; Troutwine et al., 2020). Additionally, given that the SCO receives serotonergic fiber innervation and 5-HT may regulate *sspo* expression levels (Bouchaud, 1979; Léger et al., 1983; Sakumoto et al., 1984; Richter et al., 2004), we wondered whether disruptions to the RF induced spinal curves in *tph2* mutants. To test this, we visualized the RF in *tph2* mutants using fluorescently tagged Sspo (Sspo-GFP; Troutwine et al., 2020) and followed its morphology through late larval stages, just prior to curve onset. When assessed at multiple stages between 9 and 19 dpf, we found that the RF formed normally and remained intact in *tph2* mutants (Fig 3C). No differences were detected in RF structure between mutants (*n* = 11) and control siblings (*n* = 21). This suggests RF defects are not the cause of spinal curves in *tph2* mutants.

Together, these findings argue against the hypothesis that the pineal gland is critical for maintaining spinal morphology. They further show that Tph2-dependent curvature develops despite an intact RF, thereby excluding two prominent mechanisms as explanations for spinal curvature. Instead, our findings point toward a requirement for serotonergic signaling in other neuronal populations.

### Serotonergic circuitry and locomotion are altered before scoliosis onset in *tph2* mutants

5-HT is an important neuromodulator of vertebrate spinal locomotor circuits which influence the pattern and output of locomotor activity in zebrafish (Schmidt and Jordan, 2000; Brustein et al., 2003; Montgomery et al., 2018). In larval zebrafish, a population of serotonergic ISNs arises from the ventral floor plate and develop within the spinal cord (Lillesaar et al., 2007; Montgomery et al., 2016; Chen et al., 2023). These ISNs also express *fev* (formerly *pet1*), an ETS-family transcription factor associated with serotonergic neuronal differentiation (Cheng et al., 2003; Lillesaar et al., 2007; Montgomery et al., 2016). Fev-positive (Fev+) ISNs provide a local source of spinal 5-HT, extend processes in close association with spinal motor circuitry and directly modulate spinal locomotor output (McLean and Fetcho, 2004; Montgomery et al., 2018). We therefore asked whether loss of *tph2* altered the Fev+ ISN population and whether *tph2* mutants exhibited concomitant changes in locomotor behavior.

To assess the Fev+ ISN population directly, we generated Tg(*fev:mCherryCAAX*) and Tg(*fev:mCherry-P2A-NTR2.0*) transgenic reporter lines and confirmed that mCherry+ spinal neurons co-localized with 5-HT immunoreactivity (S4A Fig). We then quantified Fev+ ISNs within the first 350 *µ*m of the rostral spinal cord at 3, 5, and 7 dpf. At all stages, *tph2* mutants exhibited significantly fewer Fev+ cells than WT controls (Fig 4A–4B; S4B Fig), indicating that *tph2* mutants contain fewer serotonergic ISNs and that this deficit is established early and persists through larval development. Complementarily, supplementing WT larvae with 5-HT produced a slight increase in Fev+ ISN number at 7 dpf (S4C Fig). Fev+ cells are distributed on both the left and right sides of the spinal cord, but we found no significant bias in their left-right distribution in either WT or *tph2* mutants (S4D Fig). Together, these findings indicate that loss of *tph2* causes an early and persistent reduction in the spinal Fev+ serotonergic population.

**Fig 4.**
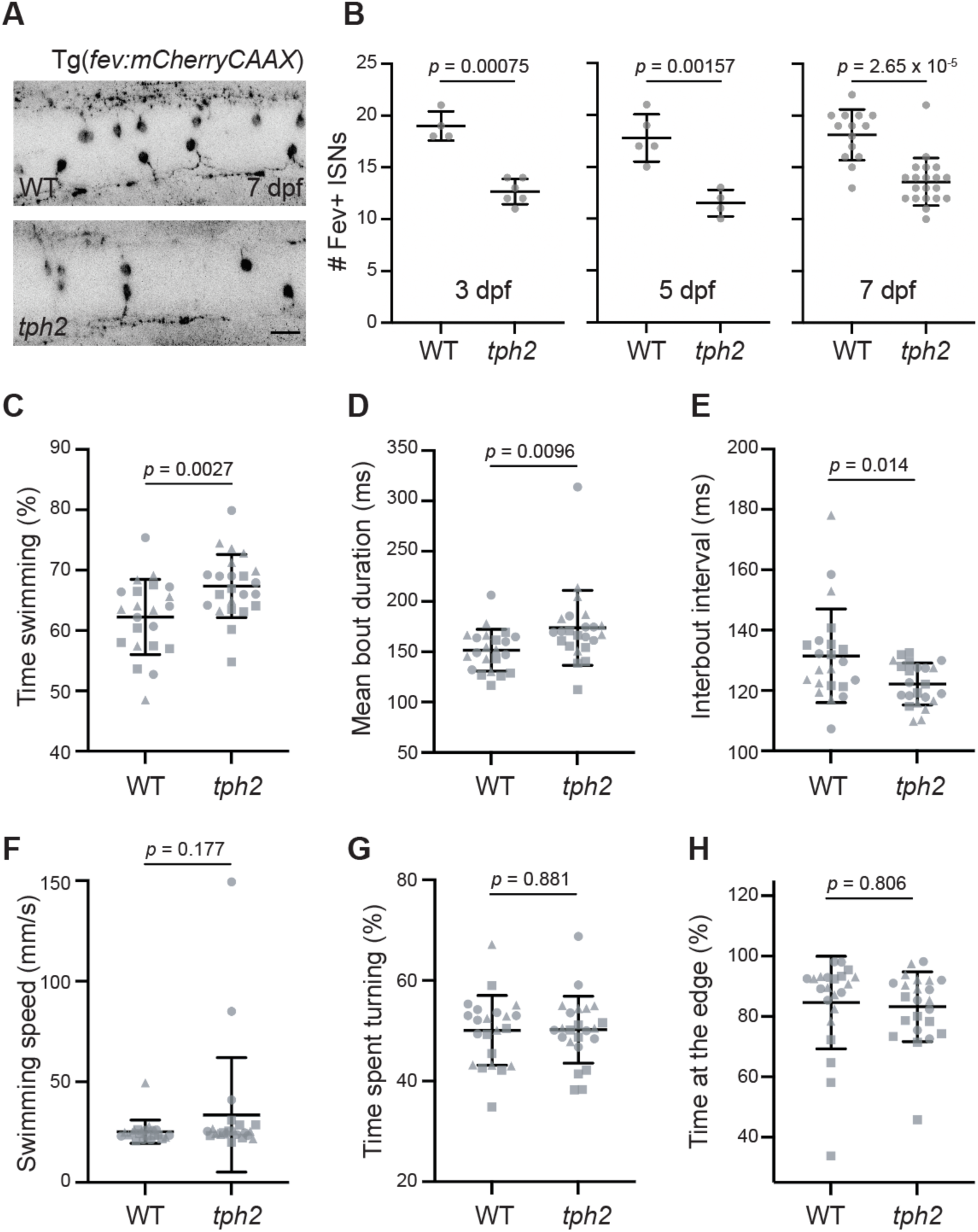
Serotonergic intraspinal neurons are reduced and locomotor behavior increased in *tph2* mutants before spinal curvature. **(A)** Representative images of Tg(*fev:mCherryCAAX*) expression in the rostral spinal cord of 7 dpf WT and *tph2* mutant larvae. Scale bar = 20 µm. **(B)** Fev+ ISNs were quantified in the first 350 *µ*m of the rostral spinal cord extending from the first Fev+ ISN. Horizontal bars show mean ± SD. Statistical comparisons were performed using two-tailed Welch’s unpaired *t* tests with Holm correction for multiple comparisons. **(C–H**) Locomotor behavior was quantified at 15 dpf before the onset of overt spinal curvature. Individual animals were recorded during free swimming and analyzed across three independent experiments, denoted by distinct symbols. Horizontal bars show mean ± SD. Statistical comparisons were performed using two-way ANOVA.

Because Fev+ ISNs are known to regulate locomotion (Montgomery et al., 2018), we next assessed *tph2* mutant locomotor behavior at 15 dpf, shortly before spinal curve onset. We recorded individual fish for 10 min in a free-swimming arena, then quantified a range of locomotion metrics (Desban et al., 2026). This revealed that *tph2* mutants spent significantly more time swimming than WT siblings (Fig 4C; *p* = 0.0027, two-way ANOVA). We asked whether this increase reflected changes in the structure of locomotor bouts, defined as discrete episodes of continuous swimming separated by interbout periods of inactivity. Mutant larvae exhibited longer swim bouts (Fig 4D; *p* = 0.0096) and shorter interbout intervals (Fig 4E; *p* = 0.014). Thus, mutants both remained active for longer and resumed swimming sooner after stopping. By contrast, average swimming speed was not significantly different between WT and mutants (Fig 4F; *p* = 0.177). This suggests that loss of *tph2* alters the temporal organization of locomotion without affecting speed.

We next asked whether the increase in locomotor activity was accompanied by broader changes in the organization of motor behavior. Turning requires coordinated asymmetric recruitment of axial musculature and provides a measure of how frequently larvae transition from forward swimming into distinct maneuvering behaviors. Because *tph2* mutants spent more time swimming overall, we normalized turning to the amount of time each fish was actively swimming. This revealed that mutants devoted the same proportion of swimming time to turning as WT larvae (Fig 4G; *p* = 0.881). Parsing data for different kinds of turns performed revealed no differences, as J-turns (*p* = 0.965), R-turns (*p* = 0.531), and C-bends (*p* = 0.463) were also unchanged between WT and mutants. Thus, despite spending more time swimming, *tph2* mutants showed no change in the frequency or composition of turning behavior.

Because serotonergic signaling is also associated with anxiety-related behavior (Maximino et al., 2013), we asked whether the increased locomotor activity in *tph2* mutants was accompanied by altered edge-seeking behavior, termed thigmotaxis (Richendrfer et al., 2012). We therefore quantified the proportion of time that larvae spent near the edge of the arena. However, *tph2* mutants did not differ from WT siblings in time spent at the edge (Fig 4H; *p* = 0.806, two-way ANOVA). This demonstrates that the locomotor phenotype is not related to changes in spatial preference within the arena. Instead, these data are consistent with a model in which loss of Tph2 causes a reduction in serotonergic ISNs, leading to increased locomotor activity. We suggest that this alters the mechanical environment experienced by the growing spine, predisposing it to abnormal curvature.

## Discussion

We found that loss of *tph2* caused a progressive three-dimensional spinal curvature that emerged during juvenile growth despite normal early vertebral patterning. Moreover, *exorh*-positive pineal photoreceptor ablation did not induce curvature, while the RF remained intact in *tph2* mutants. Instead, *tph2* mutants exhibited reduced Fev+ serotonergic ISNs and increased locomotor behavior before the onset of curvature. Our findings therefore identify serotonergic signaling as an important regulator of spinal morphology during growth and suggest a neural route through which disruption of 5-HT signaling can lead to spinal curvature.

This work has implications for our understanding of the neural control of morphology. While body morphology is often considered as an output of genetic mechanisms and developmental patterning, there is increasing evidence that neural and neuromuscular activity influences morphogenesis by controlling the mechanical environment in which tissues form and grow. For example, embryonic muscle contraction is required to maintain normal bone and joint morphology in mice (Kahn et al., 2009; Sharir et al., 2011), while in zebrafish, paralysis disrupts craniofacial skeletal and joint morphology (Shwartz et al., 2012; Brunt et al., 2016). The forces generated by the neuromuscular system, therefore, play active roles in the development of form. Similarly, by regulating locomotor activity and axial muscle behavior, our findings suggest that serotonergic circuits may influence the history of mechanical forces experienced by the growing vertebral column, thereby impacting spinal alignment. More broadly, it is becoming increasingly evident that both development and regeneration involve large-scale anatomical regulation, in which both neural and non-neural bioelectrical signaling play important roles in coordinating cell behaviors towards particular morphological outcomes (Pezzulo and Levin, 2015; Herrera-Rincon et al., 2017; Levin, 2021). In the future, it will be important to further untangle how neural activity and behavior interact with gene expression programs and tissue mechanics to shape morphology.

Related to our findings of altered locomotor behavior in *tph2* mutants, spinal alignment is also known to be influenced by improper postural and sensorimotor control. Correct posture depends on the vestibular and proprioceptive systems that drive spinal motor circuits which allow muscle activity to be continuously adjusted in response to changes in body orientation (Keshner and Cohen, 1989; Lam and Pearson, 2002; Bagnall and McLean, 2014; Delle Vedove et al., 2016; Blecher et al., 2017; Assaraf et al., 2020; Bornstein et al., 2024; Ruiz et al., 2025). Since 5-HT can impact the activity of vestibular nucleus neurons (Licata et al., 1990; Johnston et al., 1993; Han et al., 2021) and proprioceptive sensorimotor pathways (Enjin et al., 2012; Black et al., 2026), it is plausible that Tph2-associated spinal curves manifest as a result of a combination of altered locomotion, postural control, and proprioceptive feedback. Such changes could produce chronic asymmetries in mechanical loading that then produce spinal misalignment. Determining whether vestibular function and/or proprioception are altered in *tph2* mutants will be an important direction for future work.

5-HT has been previously implicated in the regulation of bone formation and remodeling. In mammals, peripheral 5-HT can directly regulate bone cells. For instance, gut-derived 5-HT has been found to inhibit osteoblast proliferation, while TPH1-dependent 5-HT can promote osteoclast differentiation (Yadav et al., 2008; Chabbi-Achengli et al., 2012). By contrast, TPH2-derived neuronal 5-HT has been proposed to regulate bone mass indirectly through central neural pathways (Yadav et al., 2009). While we cannot exclude a contribution of altered skeletal physiology to the phenotype of *tph2* mutants, we nevertheless observed normal vertebral patterning in mutants, including normal early segmentation and mineralization of the vertebral column.

Indeed, the appearance of vertebral deformation only during or after curve formation argues against a congenital defect in vertebral development as the initiating cause of spinal curvature. Our data instead suggest that the vertebral structural abnormalities in *tph2* mutants arise as a consequence of altered spinal loading. Experimental loading of growing vertebrae demonstrates that compression can limit vertebral growth, while biomechanical modeling predicts that an initial spinal curvature can generate asymmetric loading and growth, producing a self-reinforcing cycle of curve progression (Stokes et al., 1996; Stokes, 2007). In this model, an initially small deviation in alignment alters the distribution of forces across successive vertebrae, generating a positive feedback loop in which curvature promotes asymmetric growth and asymmetric growth then further amplifies the curve. A similar process is thought to contribute to AIS, where vertebral wedging and rotation are prominent components of spinal deformity, and wedging increases as curves progress in severity (Cheng et al., 2015; Labrom et al., 2020). The occasional severe abnormalities and fusions we observe in aged *tph2* mutants go beyond typical AIS pathology, however. This may reflect prolonged exposure to abnormal mechanical loading and severe deformation, rather than a direct parallel with human disease. Alternatively, species-specific differences in skeletal growth and remodeling may account for vertebral fusions that accompany otherwise AIS-like curves in *tph2* mutants.

This model is consistent with evidence that active neuromuscular control influences curve progression in AIS. Indeed, altered gait has been described in individuals with AIS (Mahaudens et al., 2009). In light of this, scoliosis exercises such as the Scientific Exercises Approach to Scoliosis (SEAS) seek to improve spinal alignment through active self-correction, postural control, and neuromotor training. Clinical studies support that these exercises can reduce curve progression in some AIS patients (Romano et al., 2015; Seleviciene et al., 2022). Our findings suggest that zebrafish could provide a useful experimental system in which to test the relationship between locomotor activity, exercise, and spinal alignment to ask whether changes in mechanical loading of the spine brought about through neuromuscular activity indeed alter the initiation, progression or recovery of spinal curvature.

Several important questions remain. First, although the altered locomotor phenotype is apparent before visible spinal curvature, our experiments do not yet demonstrate that increased locomotion causes spinal curvature. A critical next step will be to manipulate motor activity independently of Tph2. Relatedly, it will be important to directly measure axial muscle recruitment and spinal loading to determine whether locomotor changes significantly alter the mechanical forces experienced by the spine. Second, cell type-specific restoration of Tph2 will likely be a fruitful avenue for discovering the precise serotonergic neuronal populations responsible for spinal stability. Finally, it will be informative to ascertain whether the spinal column is especially vulnerable to neural and neuromuscular-associated changes during particular periods of its development, for instance during periods of rapid growth, or whether neural inputs are important for maintaining spinal morphology throughout all stages of life.

## Materials and Methods

### Zebrafish husbandry

AB and TL strains of *Danio rerio* were used. Adult zebrafish were maintained under standard laboratory conditions in a recirculating aquatic system at 28.5°C on a 14-hr light/10-hr dark cycle. Animals were maintained in an AAALAC International accredited facility. All procedures were approved by the University of Oregon Institutional Animal Care and Use Committee under protocol 21-45.

### Zebrafish lines

The transgenic and mutant zebrafish lines used in this study were Tg(*fev:mCherryCAAX*)*^b1533^*, Tg(*exorh:mCherry-P2A-NTR2.0*)*^b1516^*, Tg(*fev:mCherry-P2A-NTR2.0*)*^b1534^*, *tph2^ct817^* (Oikonomou et al., 2019), TgBAC(*entpd5a:KillerRed*)*^hu7478Tg^* (Wopat et al., 2018) and *sspo-gfp^ut24^* (Troutwine et al., 2020). Experiments were performed on zebrafish ranging from embryonic stages to 1 year of age. Sex was not determined for experiments performed at embryonic or larval stages. For juvenile and adult experiments in which sex could be determined, both male and female animals were included, and sex-specific results are reported where applicable.

### Cloning and transgenesis

The construct used to generate the Tg(*exorh:mCherry-P2A-NTR2.0*)*^b1516^* transgenic line was assembled using Gateway recombination cloning (Hartley et al., 2000; Cheo et al., 2004; Kwan et al., 2007). The mCherry-P2A-NfsB_VvF70A/F108Y coding sequence, hereafter referred to as mCherry-P2A-NTR2.0, was amplified from Addgene plasmid #158653 (Sharrock et al., 2022) using Phusion High-Fidelity DNA Polymerase (New England Biolabs) and the following primers: forward, 5′-GGGGACAAGTTTGTACAAAAAAGCAGGCTTTgccaccatggtgagcaaggg-3′; and reverse, 5′-CCACTTTGTACAAGAAAGCTGTCCCCCTAGTTttagatttcggtaaaaacag-3′ (Integrated DNA Technologies). Uppercase nucleotides indicate attB-containing adaptor sequences, and lowercase nucleotides indicate template-specific sequences. The PCR product was purified and recombined into pDONR221 using Gateway BP Clonase II Enzyme Mix (Thermo Fisher Scientific) to generate the middle entry clone pME-mCherry-P2A-NTR2.0. Multisite Gateway cloning was then performed using the pCK011 p5E-exorh promoter entry clone (Addgene plasmid #195949; Kemmler et al., 2023), pME-mCherry-P2A-NTR2.0, and p3E (Kwan et al., 2007). These entry clones were recombined into the Tol2 destination vector pDestTol2pACryGFP (Addgene plasmid #64022; Berger et al., 2013) using Gateway LR Clonase II Enzyme Mix (Thermo Fisher Scientific). The resulting *exorh* Tol2 construct was used for transgenesis.

The construct used to generate the Tg(*fev:mCherryCAAX*)*^b1533^* transgenic line was also assembled using Gateway recombination cloning. Multisite Gateway cloning was performed using the p5E-MCS-Fev promoter entry clone, pME-mCherryCAAX, and p3E (Kwan et al., 2007; Lillesaar et al., 2009). These entry clones were recombined into the Tol2 destination vector pDESTTol2pA2 using Gateway LR Clonase II Enzyme Mix (Thermo Fisher Scientific). Complete plasmid sequences were verified by whole-plasmid sequencing (Azenta Life Sciences or Plasmidsaurus).

The construct used to generate Tg(*fev:mCherry-P2A-NTR2.0*)*^b1534^* was also made using the generated middle entry clone pME-mCherry-P2A-NTR2.0. The p5E-MCS-Fev promoter entry clone, pME-mCherry-P2A-NTR2.0, and p3E (Kwan et al., 2007) were recombined into Tol2 destination vector pDESTTol2pA2 using Gateway LR Clonase II Enzyme Mix (Thermo Fisher Scientific). Complete plasmid sequences were verified by whole-plasmid sequencing (Azenta Life Sciences or Plasmidsaurus) and the resulting construct was used for transgenesis.

### Zebrafish line generation

The zebrafish lines Tg(*fev:mCherryCAAX*)*^b1533^*, Tg(*exorh:mCherry-P2A-NTR2.0*)*^b1516^* and Tg(*fev:mCherry-P2A-NTR2.0*)*^b1534^* were generated by injecting 100 ng/µL of the corresponding plasmid together with 25 ng/*µL* Tol2 transposase mRNA. Transgenic larvae were identified at 3 dpf by fluorescence in the relevant tissue and, for the *exorh* line, by the secondary eye marker. Fluorescent larvae were raised to 3 mpf and outcrossed to AB fish. F1 progeny were again screened for fluorescence, raised to 3 mpf, and outcrossed to AB fish. Before use in experiments, F2 clutches were confirmed to contain approximately 50% fluorescent embryos, consistent with transmission of a single transgene insertion. Lines were subsequently maintained by outcrossing to AB fish, with transgenic animals identified by fluorescence at 3 dpf or by genotyping from fin clips at 1 mpf or older.

### Quantitation of standard length

Zebrafish staging and standard length were determined according to Parichy et al. (2009). Fish were imaged during larval stages using a Leica S9i stereomicroscope with an integrated 10-megapixel camera. During juvenile and adult stages, fish were imaged using a mounted Google Pixel 9 Pro 50-megapixel camera. A ruler was included in each image for spatial calibration, and standard length was measured in ImageJ from the tip of the snout to the caudal peduncle. Because individual fish were not tracked longitudinally across time points, standard length was analyzed using an ordinary two-way ANOVA.

### 5-HT treatment

WT AB zebrafish embryos were exposed to exogenous 5-HT by adding 1 or 2 mM 5-HT to the rearing water from the 1-cell stage until 6 dpf. Mock controls from the same clutch were raised in parallel. 5-HT-containing water was replaced daily. At the end of the treatment, 5-HT was removed by three washes with fish water. Animals were then raised to 2.5 mpf for skeletal analysis.

### Chemical ablation

After establishment of a stable Tg(*exorh:mCherry-P2A-NTR2.0*)*^b1516^* transgenic line, transgenic fish were outcrossed to WT AB fish. One hundred 1-cell-stage embryos from the resulting clutch were transferred to embryo medium containing 1 mM metronidazole (MTZ). The working solution was prepared by diluting a 50 mM MTZ stock containing 1% DMSO 1:50 in embryo medium. At 18 hpf, embryos were dechorionated and transferred to fresh embryo medium containing 1 mM MTZ. Embryos were treated continuously with MTZ until 4 dpf and subsequently retreated for 24 hr every 3 days. Ablation of *exorh*-positive pineal photoreceptors was confirmed throughout the experiment by direct assessment of mCherry fluorescence and by immunostaining for mCherry and 5-HT.

### Faxitron imaging

X-ray images were acquired using a Faxitron UltraFocus/MultiFocus X-ray imaging system (Faxitron Bioptics). Fish were imaged either immediately after euthanasia or following fixation in 4% paraformaldehyde for up to two weeks. Fish were positioned within the imaging chamber and imaged using the instrument’s microfocus X-ray source. The voltage was set to 25 kV. Exposure time and magnification were adjusted as needed to obtain sufficient contrast and resolution for visualization of the vertebral column.

### X-ray micro-computed tomography (*µ*CT)

Whole body *µ*CT scans were acquired using a vivaCT 80 system (Scanco Medical) at 18.5 *µ*m voxel resolution for fish aged >1.5 mpf, or at 10 *µ*m voxel resolution for 1 mpf fish, as previously described (Bearce et al., 2022). High-resolution scans of individual vertebrae were acquired using a Zeiss Xradia 620 Versa µCT system at 3 *µ*m voxel resolution. Digital dissections of the spinal column were performed in 3D Slicer using the Segment Editor, and subsequent analyses were conducted as previously described (Bearce et al., 2022, 2023, Voigt et al., 2026).

### Quantitation and statistical analysis of spinal curvature

Vertebral coordinates were aligned using the approach described by Voigt et al. (2026), and spinal shape was compared between genotypes in the dorsoventral (DV) and mediolateral (ML) planes. Each fish was treated as a single biological replicate. Analyses included vertebral positions 2–24 as positions 1 and 25 were excluded because they are constrained by the alignment procedure. DV displacement was analyzed using the aligned DV coordinate (final_y), while ML curvature was analyzed using absolute ML displacement (|final_z|), as there was no evidence of a left-right bias in curve direction. The dataset contained 10 WT and 10 *tph2* mutant fish, with five males and five females of each genotype.

For whole spine comparisons, displacement at each vertebral position was modeled as a function of genotype and sex. The genotype effects across all 23 vertebral positions were squared and summed to generate a single measure of overall spinal shape difference. Significance was assessed by exact permutation of genotype labels within sex, giving 63,504 possible genotype assignments. The whole-spine statistic was recalculated for each assignment to generate the null distribution, and the permutation P value was calculated as the proportion producing a statistic at least as large as the observed value. Sex effects were assessed similarly by permuting sex within genotype. Genotype-by-sex interactions were tested by residual permutation after fitting an additive model containing both genotype and sex.

To identify individual regions of the spine that differed between genotypes, we used a max-T permutation test. At each vertebral position, displacement was modeled as a function of genotype and sex and a t statistic was calculated for the genotype effect. Genotype labels were then permuted within sex across whole fish, and the largest absolute t statistic observed across the 23 positions was retained for each permutation. These values were used to calculate multiple comparison-adjusted P values for each vertebral position.

### Immunofluorescence

Embryos were fixed overnight at 4°C in 4% paraformaldehyde with gentle rocking. Following fixation, embryos were washed in phosphate buffered saline (PBS), serially dehydrated to 100% methanol, and stored at −20°C. Before staining, embryos were rehydrated and blocked for 2 hr in blocking solution containing 5% normal sheep serum (NSS), 1% DMSO, and 0.1% Tween-20 in 1× PBS. Primary antibodies were diluted 1:500 in PBS and incubated with embryos overnight at 4°C with rocking. Embryos were then washed five times for 30 min in rinse solution containing 1% NSS, 1% DMSO, and 0.1% Tween-20 in 1× PBS.

Secondary antibodies were diluted 1:1000 in rinse solution and incubated with embryos overnight at 4°C with rocking, protected from light. Embryos were then washed five times for 30 min in rinse solution, protected from light, followed by a final 30-min wash in 0.1% Tween-20 in 1× PBS before mounting for imaging.

### Confocal microscopy

Live imaging of Sspo-GFP was performed as previously described (Bearce et al., 2022, 2025). Immunofluorescence imaging was performed using a Zeiss LSM 880 confocal microscope with 10×, 20×, and 40× water-immersion objectives and a Nikon CSU-W1 SoRa Spinning Disk confocal microscope with 4×, 10×, 20×, 40×, and 60× water-immersion objectives.

### Locomotor behavior

Locomotor behavior was assessed at 15 dpf using a behavioral recording and analysis pipeline adapted from Desban et al. (2026). Individual zebrafish were placed in shallow circular arenas (5 cm diameter, 2 mm depth) containing system water and allowed to habituate for 5 min. Fish were then recorded from above for 10 min at 25 frames per second under homogeneous illumination using an infrared imaging system.

Fish body position was tracked using ZebraZoom (Mirat et al., 2013), which identifies a series of points along the body in each video frame. Tracking data were subsequently analyzed using custom Python scripts as described in Desban et al. (2026). Fish were classified as swimming when instantaneous speed exceeded 9 mm/s, and locomotor activity was quantified as the fraction of recording time spent swimming. Individual swimming bouts were identified from transitions between moving and stationary periods, from which mean bout duration and interbout interval were calculated. Mean swimming speed was calculated from periods in which fish were actively swimming. Turning behavior was quantified from changes in body heading between successive frames. Full details are provided in Desban et al. (2026).

## Acknowledgements

We thank Judy Peirce and the Aquatic Animal Care Services Facility for zebrafish husbandry, Adam Fries and the Genomics and Cell Characterization Core, and Angela Lin and the X-Ray Imaging Core, all at the University of Oregon. We thank David Prober, Michel Bagnat, and Steve Wilson for sharing resources, Annie Fusco for technical help and Nancy-Hadley Miller for useful discussions.

## Author Contributions

Conceptualization: S.R.S., D.T.G.; Data curation: S.R.S., L.D., D.T.G.; Formal analysis: S.R.S., L.D., D.T.G.; Funding acquisition: S.R.S., J.S.E., D.T.G.; Investigation: S.R.S., B.T.B.R., M.R.S., L.D., Z.L.W.; Methodology: S.R.S., B.T.B.R., M.R.S., L.D.; Project administration: D.T.G.; Supervision: J.S.E., D.T.G.; Visualization: S.R.S., B.T.B.R., D.T.G.; Writing – original draft: S.R.S., D.T.G.; Writing – review & editing: S.R.S., J.S.E., D.T.G.

## Supplementary figures and legends

**S1 Fig.**
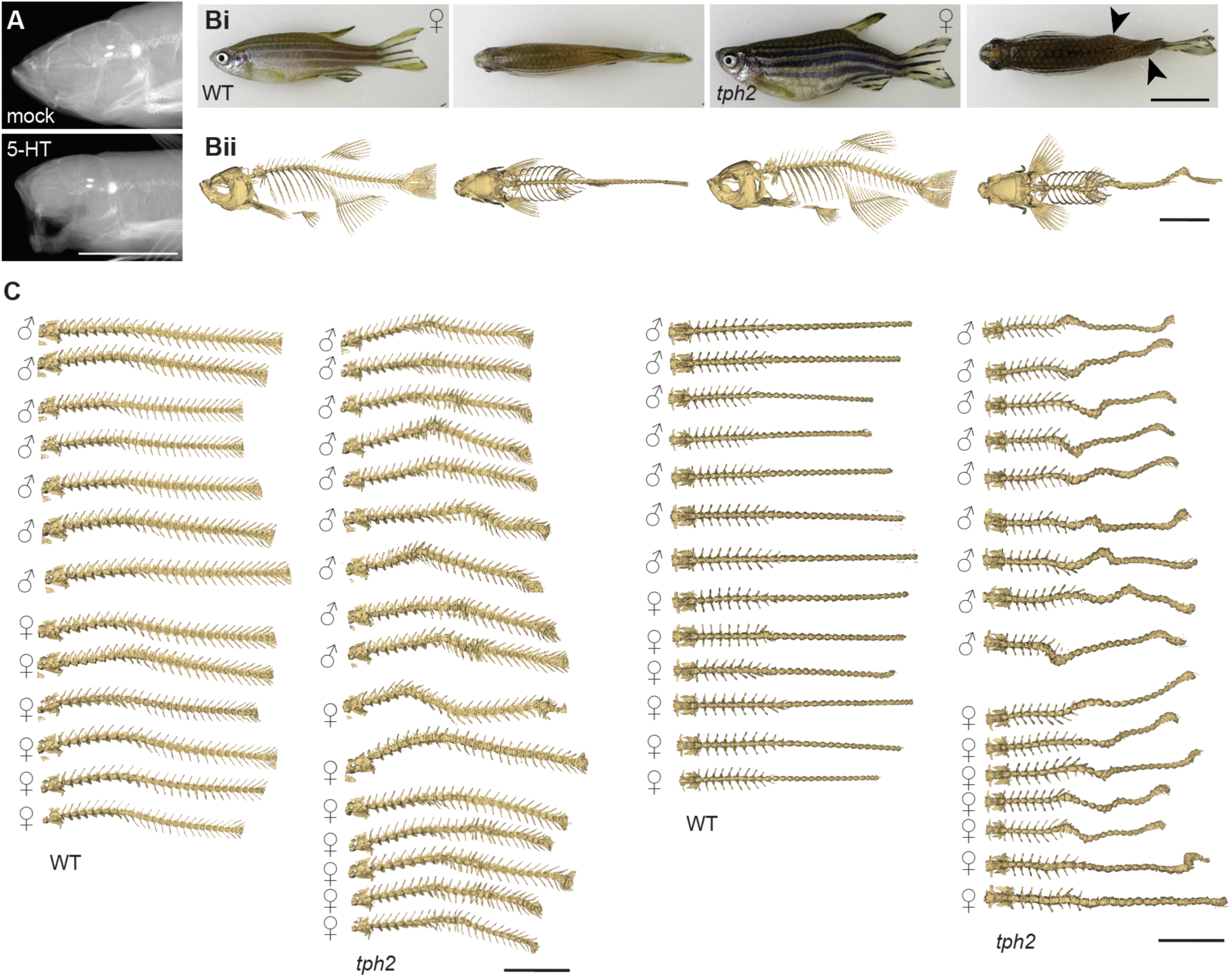
Craniofacial and spinal phenotypes associated with altered serotonergic signaling. **(A)** X-ray images of 2.5 mpf WT fish either mock-treated or treated with 2 mM 5-HT from 0–6 dpf. A proportion of treated fish developed craniofacial abnormalities (*n* = 4/14), while mock-treated fish did not (*n* = 12/12). Scale bar = 0.5 cm. **(Bi)** Lateral and dorsal views of female WT and *tph2* mutants at 6 mpf. Arrows mark the position of lateral spinal curves in mutants. Scale = 10 mm. **(Bii)** *µ*CT reconstructions of the fish in Bi. Scale bar = 10 mm. **(C)** Additional examples of *µ*CT reconstructions of 3 mpf WT and *tph2* mutant fish shown in lateral (left) and dorsal (right) views. Ribs have been removed. Scale bar = 10 mm.

**S2 Fig.**
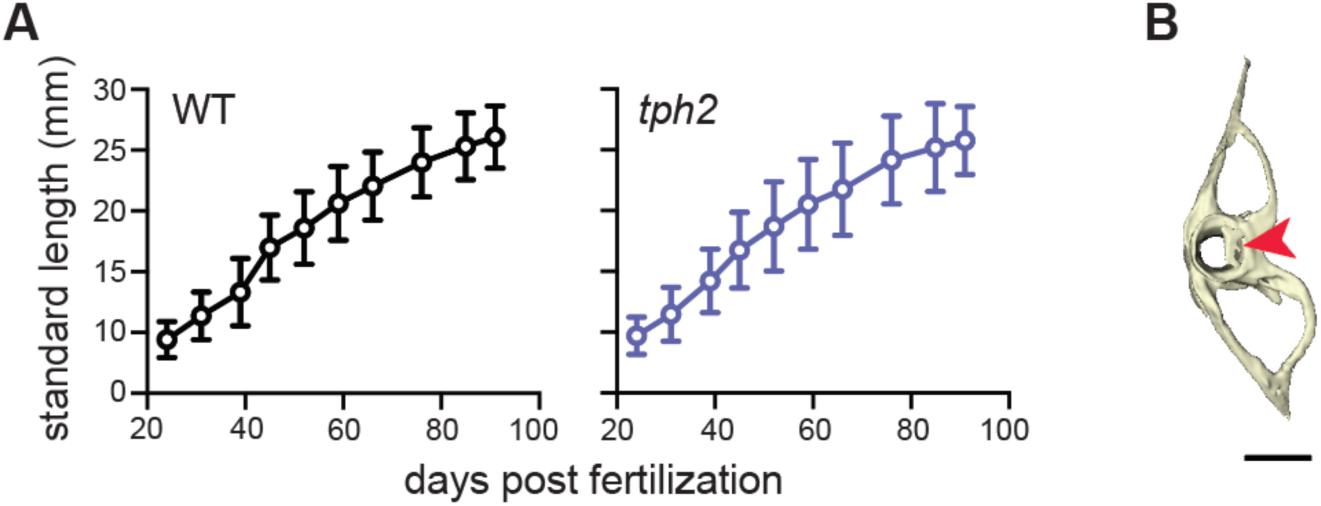
Normal growth and post-curve onset vertebral abnormalities in *tph2* mutants. **(A)** Standard length of WT and *tph2* fish during growth. Points indicate mean and error bars indicate SD. **(B)** Representative high-resolution *µ*CT reconstruction of a vertebra from a *tph2* mutant examined shortly after curve onset. Arrowhead indicates a small ectopic mineralized fragment. Scale bar = 250 *µ*m.

**S3 Fig.**
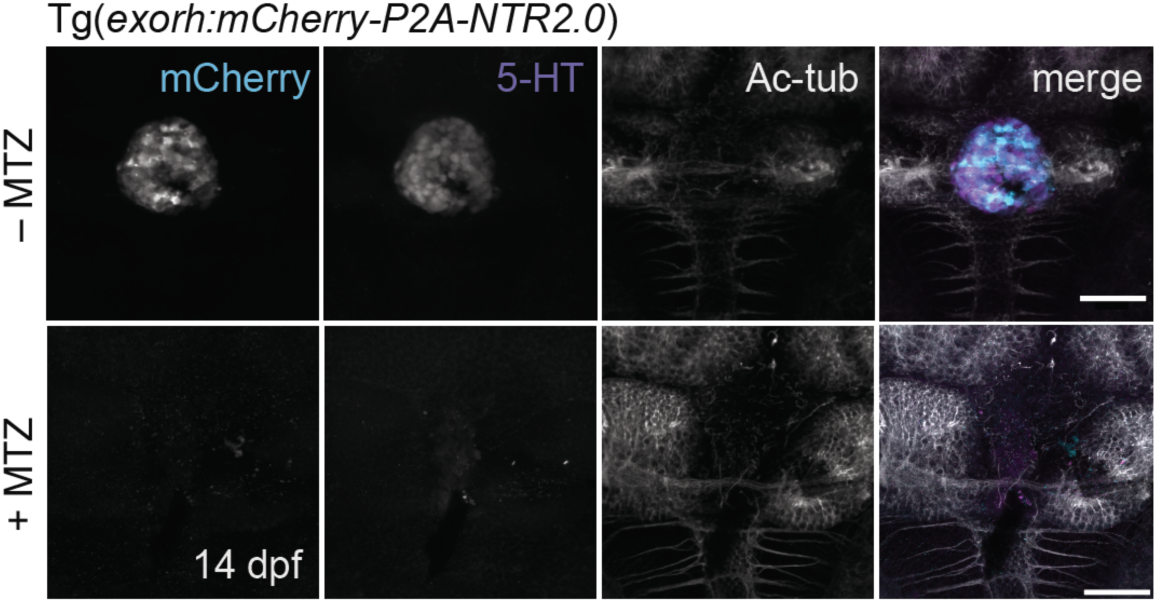
Ablation of *exorh*-positive pinealocytes following repeated MTZ treatment. Representative images of Tg(*exorh:mCherry-P2A-NTR2.0*) zebrafish at 14 dpf following repeated treatment with MTZ. Pinealocytes were visualized by mCherry fluorescence and 5-HT immunostaining. Acetylated *α*-tubulin (Ac-tub) marks surrounding neural structures. Untreated transgenic animals show robust mCherry and 5-HT signal in the pineal gland, whereas MTZ-treated animals show loss of both signals, confirming sustained pineal ablation. Scale bar = 50 *µ*m.

**S4 Fig.**
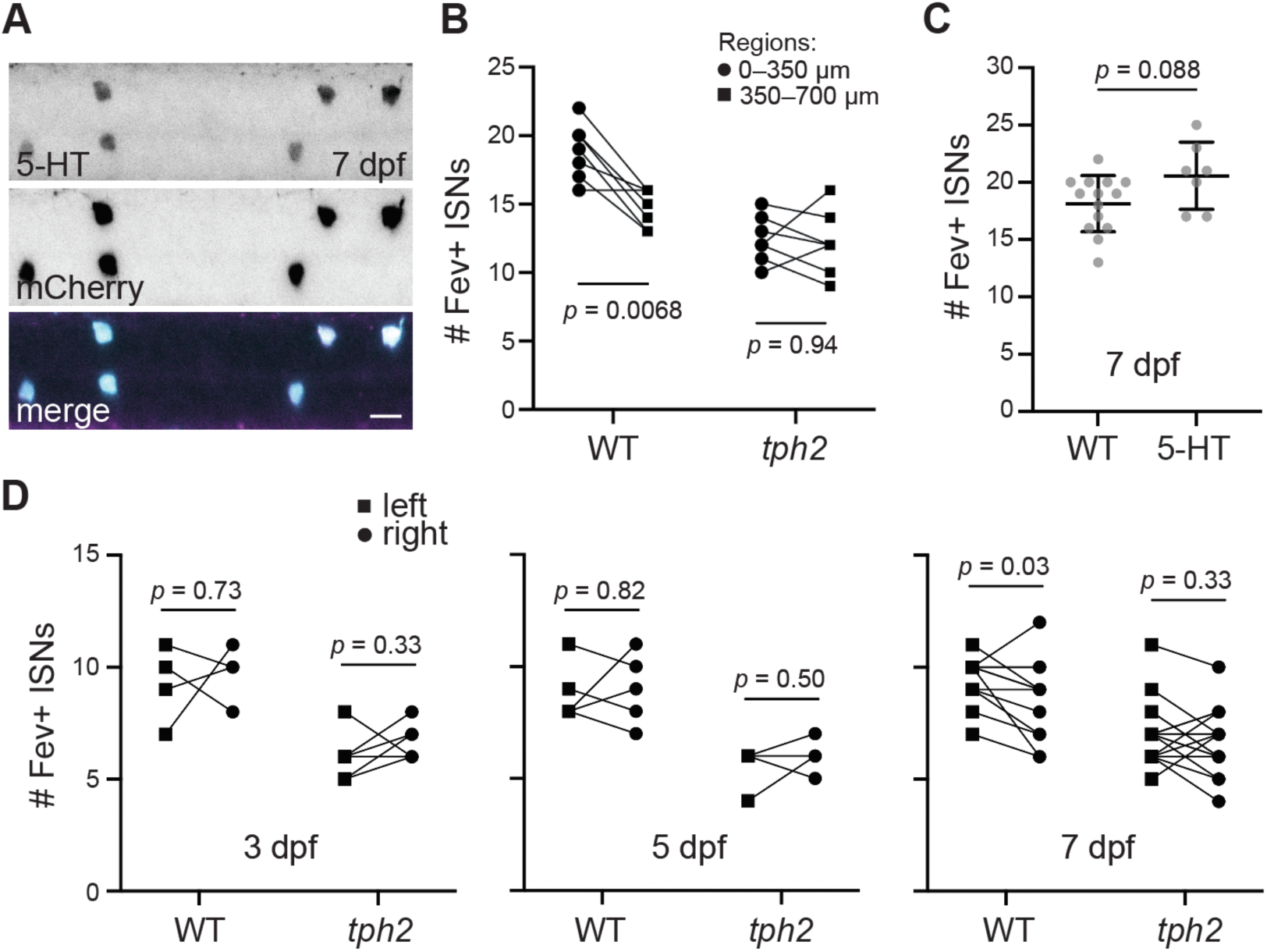
Characterization of Fev+ ISNs in *tph2* mutants and following 5-HT supplementation. **(A)** Representative image of the rostral spinal cord of a 7 dpf Tg(*fev:mCherry-P2A-NTR2.0*) transgenic larva immunostained for 5-HT and mCherry. Scale bar = 10 *µ*m. **(B)** Fev+ ISNs were quantified in two consecutive 350 *µ*m regions extending caudally from the first Fev+ ISN. Connected points represent counts from the two regions within the same animal. Two-way repeated-measures ANOVA revealed significant effects of genotype (*p* = 0.00011) and region (*p* = 0.0042). Šídák-corrected comparisons showed significantly more ISNs in the 0–350 µm than the 350–700 µm region in WT (*p* = 0.0068) but not in *tph2* mutants (*p* = 0.94). **(C)** Fev+ ISNs within the first 350 *µ*m of the rostral spinal cord, measured from the first Fev+ ISN, in WT controls (data also shown in Fig. 4B) and 2 mM 5-HT-treated fish (treated from 0–6 dpf) at 7 dpf. Horizontal bars show mean ± SD. Statistical comparisons were performed using a two-tailed Welch’s unpaired *t* test. **(D)** Fev+ ISNs within the first 350 *µ*m of the rostral spinal cord, measured from the first Fev+ ISN, plotted separately for the left and right sides. Statistical comparisons were performed using two-tailed paired *t* tests.

## Supplementary video legends

**S1 Video.** *µ*CT reconstruction of 6 mpf WT fish.

**S2 Video.** *µ*CT reconstruction of 6 mpf *tph2* mutant fish.

**S3 Video.** *µ*CT reconstruction of two-vertebra segment from a 12 mpf WT fish.

**S4 Video.** *µ*CT reconstruction of two-vertebra segment from a 12 mpf *tph2* mutant showing vertebral fusion.

**S5 Video.** *µ*CT reconstruction of two-vertebra segment from a 28 dpf *tph2* mutant showing vertebral warping.

## Abbreviations

5-HT: 5-hydroxytryptamine
AIS: adolescent idiopathic scoliosis
CNS: central nervous system
CSF: cerebrospinal fluid
dpf: days post fertilization
DV: dorsoventral
ISN: intraspinal neuron
*µ*CT: micro-computed tomography
ML: mediolateral
mpf: months post fertilization
MTZ: metronidazole
NTR: nitroreductase
RF: Reissner fiber
SCO: subcommissural organ
SEAS: Scientific Exercises Approach to Scoliosis
Sspo: SCO-spondin
Tph2: tryptophan hydroxylase 2
WT: wild type

